# Role of XerCD in release of over-replicated DNA through Outer Membrane Vesicles in *Escherichia coli*

**DOI:** 10.1101/2023.05.30.542921

**Authors:** Johannes Mansky, Hui Wang, Irene Wagner-Döbler, Jürgen Tomasch

**Affiliations:** Institute of Microbiology, Technical University of Braunschweig, Braunschweig, Germany; Laboratory of Anoxygenic Phototrophs, Institute of Microbiology of the Czech Academy of Science – Centre Algatech, Třeboň, Czech Republic

## Abstract

Outer membrane vesicles (OMVs) are universally produced by prokaryotes and play important roles in symbiotic and pathogenic interactions. Here we show that the Gammaproteobacterium *Escherichia coli* produces OMVs that contain DNA enriched for the region around the terminus of replication *ter*, and specifically for the recognition sequence *dif* of the two site-specific recombinases XerCD, similar to OMVs from the Alphaproteobacterium *Dinoroseobacter shibae*. In deletion mutants of *xerC* or *xerD*, the enriched region around *ter* becomes broader while the peak directly at the *dif* sequence itself is reduced.

**Importance:** Imprecise termination of replication can lead to over-replicated parts of bacterial chromosomes that have to be excised and removed from the dividing cell. The underlying mechanism is so far poorly understood. Our data suggest a conserved mechanism for repair and removal of over-replicated DNA through outer membrane vesicles and an active role of the site-specific XerCD recombinase complex therein.

## Introduction

Membrane vesicles are excreted by cells from all domains of life, and their cargo and the physiological roles discovered until now are as diverse as life itself [1][2]. Outer membrane vesicles of Gram negative bacteria (OMV) have often been found to contain DNA, but it is unclear how the DNA could have been transferred from the cytosol to the periplasmic space and into the vesicles, unless as a result of explosive cell lysis [3].

We had previously shown that the Alphaproteobacterium *Dinoroseobacter shibae* produces OMVs constitutively during growth [4]. Based on multiple evidence we hypothesized that these vesicles, which were ejected from the dividing cell’s division plane, act as a means to remove over-replicated chromosomal DNA at the end of the cell cycle. The key observation behind this hypothesis was the enrichment of DNA from the terminus of replication *ter*, and specifically the *dif* site located in the center of *ter*, in the vesicle lumen [4]. The latter is a 28 bp palindromic sequence which is recognized by two site-specific recombinases XerC and XerD [5][6]. When both replication forks of circular chromosomes meet at *ter*, they collide with the divisome complex, and the XerC/XerD enzymes are activated by FtsK to resolve chromosome dimers, resulting from illegitimate recombination between left and right replichore in a fraction of the population and lethal for the cells [7]. The two replication forks often do not collide exactly at *ter*, because the left and right replichores can progress with different speed, resulting in over-replication of DNA - including *dif* - around *ter* [8][9][10][11]. The DNA enriched in OMV might therefore originate from over-replication repair. It had to remain open if the XerCD enzymes themselves are active in this repair process, which would imply that they have a second role beyond dimer resolution, or if other enzymes are involved as well [9].

To test our hypothesis further, here we studied two questions: (1) Is the enrichment of the *dif* site specific for *D. shibae*, an Alphaproteobacterium from the Roseobacter group, or does it occur in other Proteobacteria as well? We chose *Escherichia coli* as a second model because it is the archetypical, best understood organism regarding the mechanisms of replication and cell division [12][13] and a library of well-characterized clean gene knockouts including the two genes of interest is available [14]. (2) Are the XerCD enzymes directly involved in the excision of over-replicated DNA fragments around *ter*, implicating that they have a second function in addition to resolving chromosome dimers? Therefore, we investigated the DNA composition in the lumen of OMVs produced by deletion mutants of XerC and XerD in *E. coli*.

## Methods

Strains *E. coli* K-12 BW251113 (WT), *E. coli* JW3784 (Δ*xerC*) and *E. coli* JW2862 (Δ*xer*D) were obtained from the Keio Collection [14]. They were grown on LB plates or liquid LB medium at 37°C, liquid cultures with shaking at 180 rpm. Cell count was determined by flow cytometry using a MacsQuant Analyzer 10, and vesicle count was determined using the NanoSight NS300 (Malvern Panalytical). Vesicles were purified from 1 L of culture per replicate using a tangential flow filtration system (Vivaflow 200, Sartorius). Details on all methods, including DNA isolation, can be found in [4]. Libraries for sequencing were prepared with NEBnext Ultra IIFS DNA kit according to the manufacturer’s protocol. 50 bp paired-end sequencing was performed on the Novaseq 6000 to a depth of 2 million reads per sample. Quality trimming of raw reads was conducted with sickle v.1.33. Processing and analysis of sequencing data were performed as described [4]. Sequence reads for two to three replicate samples were deposited at the European Nucleotide Archive (ENA; https://www.ebi.ac.uk/ena) under accession number <u>PRJEB62439</u>.

### Influence of *xerC* and *xerD* knock outs on the DNA content of *E. coli* OMVs

The Δ*xerC* and Δ*xerD* mutants grew at the same rate as the wild-type (Figure 1A), thus they do not have an obvious fitness defect, in accordance with the published strain descriptions [14]. However, the dynamics of OMVs production was different in the mutants. While the OMV concentration in the supernatant remained stable around 2*10^8^ vesicles/ml for the wild type, it increased from a similar initial value to 5*10^9^ vesicles/ml for the mutants during the 12 hours of cultivation (Figure 1B). The ratio of vesicles to cells decreased for the wild-type from 10:1 to 0.3:1 while it remained between 3:1 and 10:1 for both mutants (Figure 1C). If our hypothesis is true and the DNA in OMVs represents excised over-replicated fragments, then the mutants apparently had more waste to get rid of.

**Figure 1.**
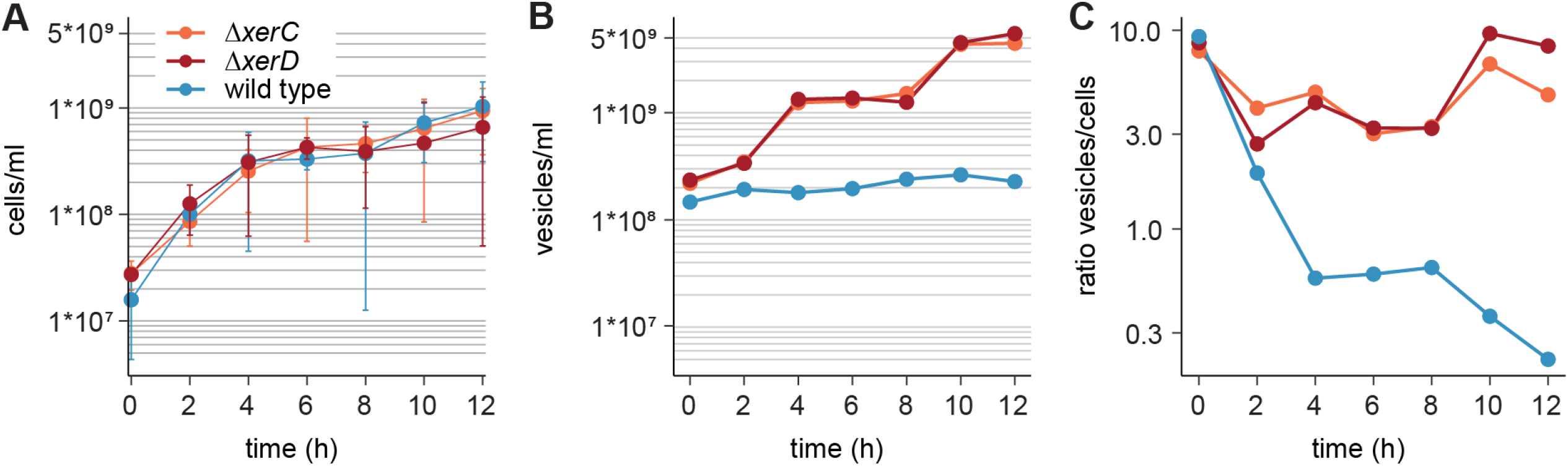
Growth and outer membrane vesicle production of *E. coli*. **(A)** Growth of E. coli wild type and mutant strains. **(B)** Vesicles in the supernatant of E. coli strains during growth. **(C)** Ratio of vesicles to cells during growth.

Sequencing of the DNA isolated from the OMV’s lumen clearly showed that a 100 kb region around *ter* and particularly the *dif* sequence almost in the center was enriched more than 120 fold compared to the rest of the chromosome (Figure 2A). This is comparable to the 200 kb region surrounding the homologues site and also found up to 120 fold enriched in OMVs of *D. shibae* [4]. Given the large phylogenetic distance between those two organisms, this observation suggests that a conserved mechanism might govern the DNA composition of this type of OMVs.

**Figure 2.**
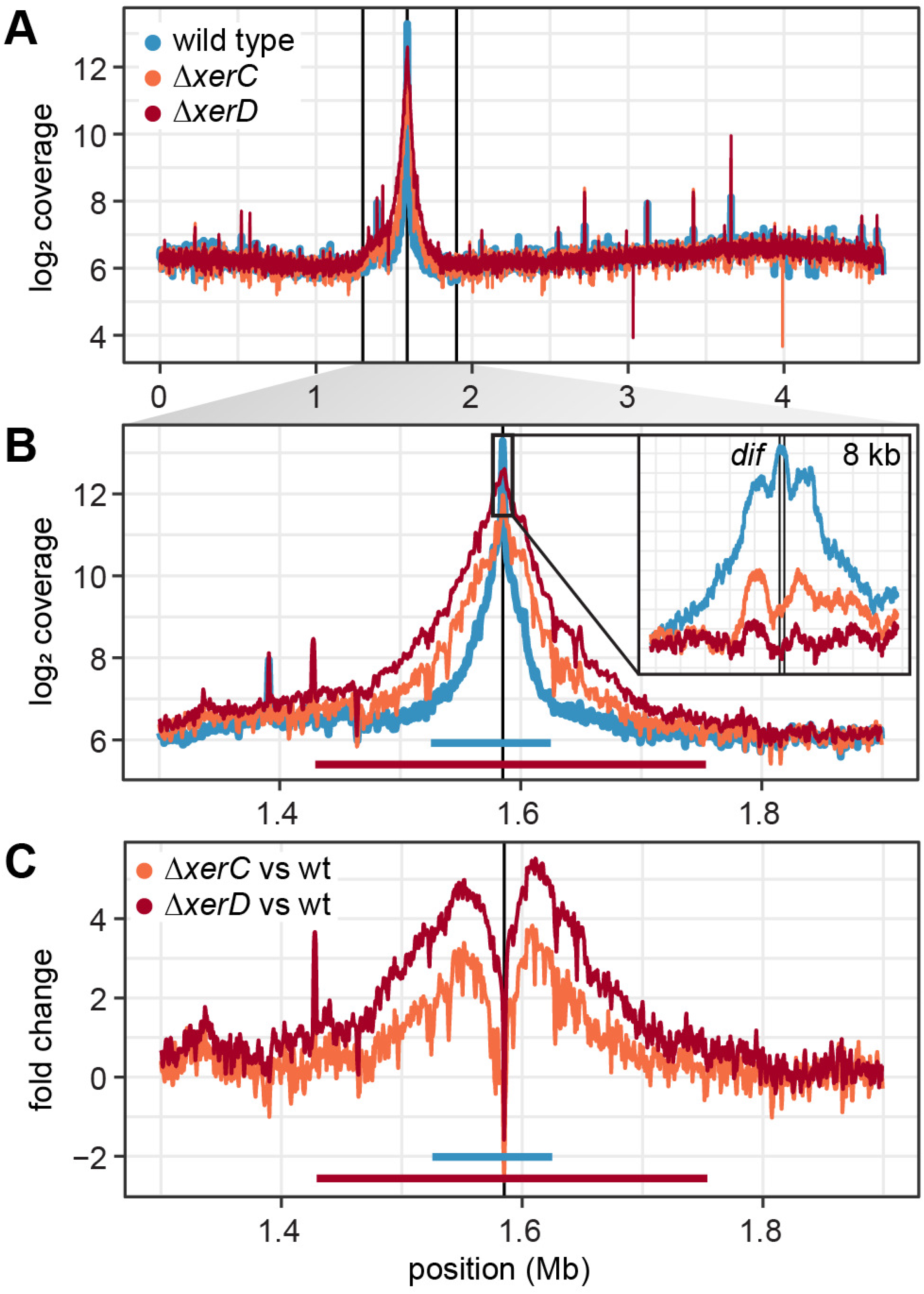
DNA content of *E. coli* outer membrane vesicles. **(A)** Coverage of mapped reads on the chromosome of *E. coli* averaged for sliding windows of 0.5 kb **(B)** Zoom in to the *ter* region. The peak ranges for the wild-type and mutants are marked. The inset shows the *dif* site with single nucleotide resolution. **(C)** Fold change between the coverage of the *ter* region in the mutants compared to the wild-type.

The enrichment of the *ter* region in DNA of OMVs from either mutant clearly differed from that of the wild-type (Figure 2B). The peak range increased asymmetrically to approximately 350 kb around *dif* with this broader region being up to four fold higher present in the mutant OMVs, suggesting increased and lengthened over-replication in these strains (Figure 2B and C). A single-nucleotide-view on the most strongly enriched region revealed three peaks with the central maximum at the 28 bp *dif* site for the wild type (Figure 2B). This maximum was 2.5-fold reduced in both mutants, while the surrounding two peaks were still visible, particularly in Δ*xerC*. These data suggest impaired binding or functioning of the XerCD recombinase complex at the *dif* site when one of the homologs is knocked out.

### Roles of XerCD recombinase in over-replication repair

The peak of DNA in OMVs at *dif* suggests that this site acts as an anchoring point for over-replication repair. The increased length of excised DNA fragments and the reduced representation of *dif* itself in the mutants suggest the site-specific recombinases XerCD must therein play an active role which is not yet understood. The XerCD recombination complex is highly adaptable. It was used for the construction of markerless gene deletions [15][16]. It is exploited by phages and plasmids for integration into the chromosome [5][17], and in some bacteria only one recombinase is required [18]. Both XerC and XerD can efficiently mediate recombination independently as shown by reporter plasmids carrying tandem *dif* sites [19]. We hypothesize that in the functionally impaired Δ*xerC* and Δ*xerD* mutants over-replication has become more likely and is to a lesser extent resolved directly at *dif*. Possible mechanisms might involve a delayed recruitment of the chromosome segregation machinery to *dif* [20] or impaired interaction with either the RecBCD enzymes required for excision of over-replicated regions [9] or the Tus proteins acting as barriers against over-replication [21].

To conclude, we show that the enrichment of the *ter* region of the bacterial chromosome in OMVs it not restricted to *D. shibae*, but also found in *E. coli*. The site-specific recombinases XerC and XerD play an active role for enrichment of their recognition sequence *dif* in the lumen of OMVs. Given their almost universal presence in prokaryotes [22] and the extreme conservation of the cell division molecular machinery it would be interesting to unravel the underlying mechanisms in more detail.

## References

1. Gill S, R Catchpole, P Forterre. Extracellular membrane vesicles in the three domains of life and beyond. FEMS Microbiol Rev 2019; 43: 273–303.

2. Toyofuku M, S Schild, M Kaparakis-Liaskos, L Eberl. Composition and functions of bacterial membrane vesicles. Nat Rev Microbiol 2023.

3. Cárcamo-Oyarce G, LG Monahan, IG Charles, N Nomura, L Turnbull, X Yap, S Ito, R Shimoni, M Kurosawa, AL Hynen, SR Osvath, CB Whitchurch, U Omasits, M Toyofuku, R Cavaliere, CH Ahrens, L Eberl, ES Gloag, G Pessi, et al. Explosive cell lysis as a mechanism for the biogenesis of bacterial membrane vesicles and biofilms. Nat Commun 2016; 7.

4. Wang H, N Beier, C Boedeker, H Sztajer, P Henke, M Neumann-Schaal, J Mansky, M Rohde, J Overmann, J Petersen, F Klawonn, M Kucklick, S Engelmann, J Tomasch, I Wagner-Döbler. Dinoroseobacter shibae outer membrane vesicles are enriched for the chromosome dimer resolution site dif. mSystems 2021; 6.

5. Castillo F, A Benmohamed, G Szatmari. Xer site specific recombination: Double and single recombinase systems. Front Microbiol 2017; 8: 1–18.

6. Midonet C, F-X Barre. Xer Site-Specific Recombination: Promoting Vertical and Horizontal Transmission of Genetic Information. Microbiol Spectr 2014; 2: 1–18.

7. Keller AN, Y Xin, S Boer, J Reinhardt, R Baker, LK Arciszewska, PJ Lewis, DJ Sherratt, J Löwe, I Grainge. Activation of Xer-recombination at dif: Structural basis of the FtsKγ-XerD interaction. Sci Rep 2016; 6: 1–12.

8. de Septenville AL, S Duigou, H Boubakri, B Michel. Replication fork reversal after replication-transcription collision. PLoS Genet 2012.

9. Wendel BM, CT Courcelle, J Courcelle. Completion of DNA replication in Escherichia coli. Proc Natl Acad Sci U S A 2014.

10. Hiasa H, KJ Marians. Tus prevents overreplication of oriC plasmid DNA. J Biol Chem 1994.

11. Lloyd RG, CJ Rudolph. 25 years on and no end in sight: a perspective on the role of RecG protein. Curr Genet 2016; 62: 827–840.

12. Du S, J Lutkenhaus. Assembly and activation of the Escherichia coli divisome. Mol Microbiol 2017; 105: 177–187.

13. Graumann PL. Chromosome architecture and segregation in prokaryotic cells. J Mol Microbiol Biotechnol 2014; 24: 291–300.

14. Baba T, T Ara, M Hasegawa, Y Takai, Y Okumura, M Baba, KA Datsenko, M Tomita, BL Wanner, H Mori. Construction of Escherichia coli K-12 in-frame, single-gene knockout mutants: The Keio collection. Mol Syst Biol 2006; 2.

15. Cranenburgh RM, AE Bloor. An efficient method of selectable marker gene excision by Xer recombination for gene replacement in bacterial chromosomes. Appl Environ Microbiol 2006; 72: 2520–2525.

16. Debowski AW, JC Gauntlett, H Li, T Liao, M Sehnal, HO Nilsson, BJ Marshall, M Benghezal. Xer-cise in Helicobacter pylori: One-step Transformation for the Construction of Markerless Gene Deletions. Helicobacter 2012; 17: 435–443.

17. Fournes F, E Crozat, C Pages, C Tardin, L Salomé, F Cornet, P Rousseau. FtsK translocation permits discrimination between an endogenous and an imported Xer/dif recombination complex. Proc Natl Acad Sci U S A 2016; 113: 7882–7887.

18. Nolivos S, C Pages, P Rousseau, P Le Bourgeois, F Cornet. Are two better than one? Analysis of an FtsK/Xer recombination system that uses a single recombinase. Nucleic Acids Res 2010; 38: 6477–6489.

19. Grainge I, C Lesterlin, DJ Sherratt. Activation of XerCD-dif recombination by the FtsK DNA translocase. Nucleic Acids Res 2011; 39: 5140–5148.

20. Bhowmik BK, AL Clevenger, H Zhao, V V. Rybenkov. Segregation but Not Replication of the Pseudomonas aeruginosa Chromosome Terminates at Dif. MBio 2018; 9: 1–13.

21. Mulcair MD, PM Schaeffer, AJ Oakley, HF Cross, C Neylon, TM Hill, NE Dixon. A Molecular Mousetrap Determines Polarity of Termination of DNA Replication in E. coli. Cell 2006; 125: 1309–1319.

22. Kono N, K Arakawa, M Tomita. Comprehensive prediction of chromosome dimer resolution sites in bacterial genomes. BMC Genomics 2011; 12.

